# Do the Moroccan SARS-CoV-2 genetic diversity hamper the use of the developed universal vaccines in Morocco?

**DOI:** 10.1101/2020.06.30.181123

**Authors:** Meriem Laamarti, Abdelomunim Essabbar, Tarek Alouane, Souad Kartti, Nasma Boumajdi, Houda Bendani, Rokia Laamarti, Loubna Allam, Mouna Ouadghiri, M.W. Chemao-Elfihri, Fatima Ghrifi, Imane Smyej, Jalila Rahoui, Houda Benrahma, Idrissa Diawara, Tarik Aanniz, Naima El Hafidi, Rachid El Jaoudi, Chakib Nejjari, Saaid Amzazi, Rachid Mentag, Lahcen Belyamani, Azeddine Ibrahimi

## Abstract

The SARS-CoV-2 identified as coronavirus species associated with severe acute respiratory syndrome. At the time of writing, the genetic diversity of Moroccan strains of SARS-CoV-2 is poorly documented. The present study aims to analyze and identify the genetic variants of forty-eight Moroccan strains of SARS-CoV-2 collected from mid-March to the end of May and the prediction of their possible sources. Our results revealed 108 mutations in Moroccan SARS-CoV-2, 50% were non-synonymous were present in seven genes (S, M, N, E, ORF1ab, ORF3a, and ORF8) with variable frequencies. Remarkably, eight non-synonymous mutations were predicted to have a deleterious effect for (ORF1ab, ORF3a, and the N protein. The analysis of the haplotype network of Moroccan strains suggests different sources of SARS-CoV-2 infection in Morocco. Likewise, the phylogenetic analysis revealed that these Moroccan strains were closely related to those belonging to the five continents, indicating no specific strain dominating in Morocco. These findings have the potential to lead to new comprehensive investigations combining genomic data, epidemiological information, and clinical characteristics of SARS-CoV-2 patients in Morocco and could indicate that the developed vaccines are likely to be effective against Moroccan strains.

## Introduction

The novel coronavirus 2019, also known as severe acute respiratory syndrome Coronavirus 2 (SARS-CoV-2) (1), is the causative agent of coronavirus disease-2019 (COVID-19), a new type of pneumonia that has caused an epidemic in Wuhan, China, in late December 2020. The virus’s fast transmission worldwide and a large number of confirmed cases made the World Health Organization (WHO) declare COVID-19 as a global pandemic on March 11, 2020 (2). As of October 11, 2020, the virus has spread to 235 different countries, infected more than 37 million people, and caused more than one million deaths (https://covid19.who.int/). It should be noted that mortality from SARS-CoV-2 differs considerably by geographic region. In Morocco, the first case of COVID-19 was identified on March 02. Since then, the number of infections and the number of deaths has been increasing continuously; by October 11, 2020, the Moroccan ministry of Health announced 149,841 confirmed cases, including 2,572 deaths.

SARS-CoV-2 is a positive-sense single-stranded RNA virus, encoding four structural proteins (spike (S), envelope (E), membrane (M) and nucleocapsid (N), 16 non-structural proteins (nsp1 to nsp16), and five accessory proteins (ORF3a, ORF6, ORF7a, ORF7b, and ORF8) (3, 4). Among these, two genes are considered to be the most important targets for candidate vaccines: the S protein, which is responsible for the binding to host cells membrane receptors (ACE2) via its receptor-binding domain (RBD), and the RNA-dependent RNA polymerase (RdRp, also called nsp12) which is a key part of the virus replication/transcription machinery (5-7).

It is known that the mutation rate of the RNA virus contributes to viral adaptation, creating a balance between the integrity of genetic information and the variability of the genome, thus allowing viruses to evade the host’s immune system and develop drug resistance (8, 9). Indeed, recent studies have reported specific genotypes at particular geographic locations and that the evolution of SARS-CoV-2 over time shows a mechanism of co-accumulation of mutations that could potentially affect the spread and the severity of this virus (OYW).

At the time of writing, genetic variants, their impact, and distribution along the viral genome of Moroccan strains are poorly documented. Few studies investigating the genetic characteristics have been done so far. The analysis of 20 Moroccan SARS-CoV-2 genomes suggested that the epidemic spread in Morocco did not show a predominant SARSCoV-2 route and took origin from multiple and unrelated sources (8). In this study, we investigated the level of diversity of 48 strains of SARS-CoV-2 that were collected in Morocco between mid-March and the end of May 2020, including nine new strains. The potential source of these strains was also predicted by comparison to other genomes from six continents.

## Materials and Methods

### Data collection and Genomes sequencing

Thirty-nine Moroccan genomes were collected from the GISAID database (http://gisaid.org) (10). Also, nine genomes were sequenced in-house recently to build a dataset counting 48 Moroccan genomes.

The nine genomes’ sequencing was achieved by the routine workflow on the MinION Mk1B Nanopore platform using the R9.4.1 flowcell. The viral RNA was extracted from nine clinical samples, and the cDNA was synthesized using a kit (Roche) with random hexamers. Genome enrichment was done by Q5 Hot Start High-Fidelity DNA Polymerase (NEB), following the manufacturer’s specifications, using a set of primers designed by the ARTIC network (https://artic.network/ncov-2019) that target overlapping regions of the SARS-CoV-2 genome. The PCR products were purified by adding an equal volume of AMPure XP beads (Beckman Coulter). The sequencing was performed according to the eight-hour routine workflow, and amplicons were repaired with NEBNext FFPE Repair Mix (NEB), followed by the DNA ends preparation using NEBNext End repair/ dA-tailing Module (NEB) before adding native barcodes and sequencing adapters supplied in the EXP-NBD104/114 kit (Nanopore) to the DNA ends. After priming the flow cell, 60 ng DNA per sample was pooled with a final volume of 65 uL. Following the ligation sequencing kit (SQK-LSK109) protocol, the sequencing was performed using the MinION Mk1B device.

### The assembly

The gupplyplex and minion scripts of the ARTIC Network bioinformatics protocol (https://artic.network/ncov-2019/ncov2019-bioinformatics-sop.html) were used for reads Preprocessing and Consensus Building for nanopore sequencing. Gupplyplex (default parameters) was used for quality control and filtering of reads (--min-length 400 --max-length 700) followed by MinION pipeline (default parameters) to perform the sequences mapping, primers trimming, variations calling, and consensus assembly building.

### Variant calling analysis

A set of 39 SARS-CoV-2 genomes were downloaded from the GISAID database (http://www.gisaid.org/) and added to the nine newly sequenced genomes (10)(**Table 1**).

The reads generated by MinION Nanopore-Oxford of the nine isolates were mapped to the reference sequence genome Wuhan-Hu-1/2019 using BWA-MEM v0.7.17-r1188 (11) with default parameters, while the data downloaded from the GISAID database were mapped using Minimap v2.12-r847 (12)

The BAM files were sorted using SAMtools (13)and were subsequently used to call the genetic variants in variant call format (VCF) by BCFtools (13). The final call set of the 48 genomes was annotated, and their impact was predicted using SnpEff v 4.3t (14). First, the SnpEff databases were built locally using annotations of the reference genome NC_045512.2 obtained in the GFF format from the NCBI database. Then, the SnpEff database was used to annotate SNPs and with putative functional effects according to the categories defined in the SnpEff manual (http://snpeff.sourceforge.net/SnpEff_manual.html). The variants were also evaluated for functional consequences using the SIFT algorithm (34). The SIFT prediction is given as a Tolerance Index score ranging from 0.0 to 1.0, which is the normalized probability that the amino acid change is tolerated. SIFT scores less than or equal to 0.05 are predicted by the algorithm to be intolerant or deleterious amino acid substitutions, while scores greater than 0.05 are considered tolerant.

### Phylogenetic and haplotype network analysis

In order to determine the source(s) of strains circulating in morocco, we performed a multiple sequence alignment using Muscle v 3.8 (15) for the 48 Moroccan strains with 225 genomes of SARS-CoV-2 from Africa, Asia, Europe, North, South America, and Oceania (**Supplementary Material; Table S2**). Maximum-likelihood trees were inferred with IQ-TREE v1.5.5 under the GTR model (16). Generated trees were visualized *using FigTree* 1.4.3 To generate haplotypes for the 48 Moroccan genomes, we aligned the viruses’ complete genomes using Muscle v 3.8 (15).

The generated (.fasta) file was converted to a (.meg) file using a home-made script. Using the dnasp5 tool (17), we generated the file (.rdf), compatible with the NETWORK Package, which allowed the network’s tracing linking the 48 genomes’ haplotypes, using the MJ (Median Joining). In order to estimate genealogical relationships of haplotype groups, the phylogenetic networks were inferred by PopART package v1.7.2 (18)using the TCS method and MSN, respectively.

## Results

### Genetic diversity of Moroccan strains of SARS-CoV-2

To identify and study the genetic variants of the SARS-COV-2 genomes from Morocco, 48 genomes were collected from mid-March to the end of May; including nine strains sequenced in the present study (**Supplementary Material, Table S1**). 94.9% to 99.93% of the reads produced for the nine genomes were correctly mapped to the Wuhan-Hu-1/2019 reference sequence. Analysis of genetic variants revealed a total of 108 variant sites in the 48 genomes analyzed (**Figure 1**); including 54 (50%) non-synonymous, 42 (38.89%) synonymous, and 3 frame-shifts (2.78%). The remaining (8.33%) were distributed along the intergenic regions.

**Figure 1:**
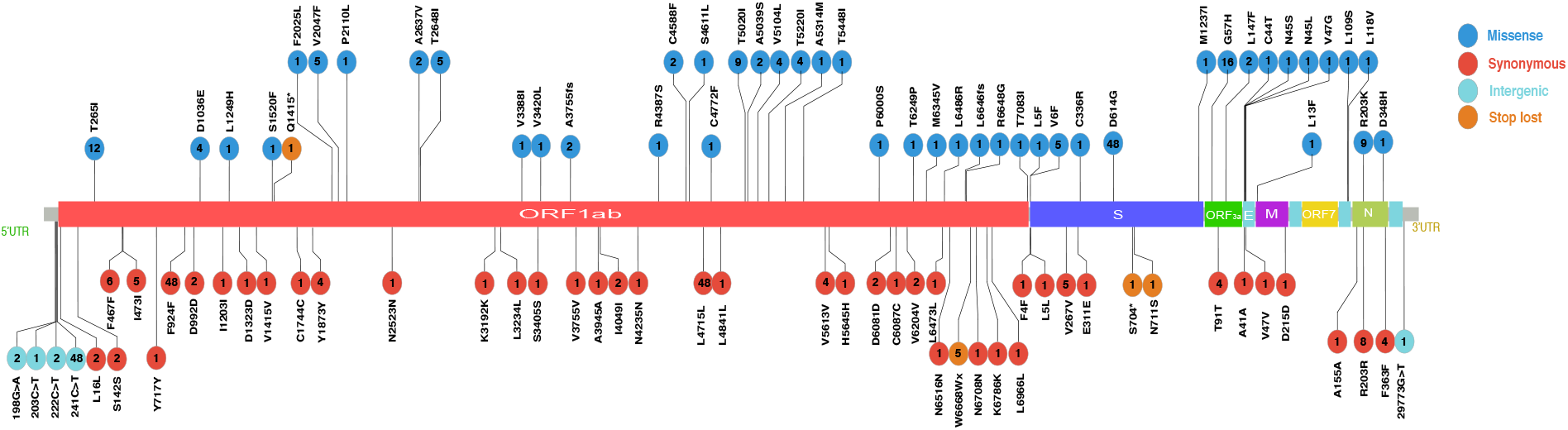
SARS-CoV-2 genomes landscape illustration representing mutations identified in 48 Moroccan genomes. Colored circles represent gene distribution across the genomes. Lollipop stick represents individual mutation. Colored circles represent the type of mutation and the blue box represents the number of genomes harboring the mutations.

Interestingly, 36.11% of the total mutations were shared between at least two genomes, while the rest were singleton mutations (unique to one genome). Mutational distribution along the viral genome revealed seven affected genes (ORF1ab, S, N, M, E, ORF8, and ORF3a) with varying mutational frequencies. Among the non-synonymous mutations, 28 were located in the ORF1ab gene and were present in eight non-structural proteins; 9 mutations in the nsp12-RNA-dependent RNA polymerase (RdRp) (C4588F, S4611L, C4772F, T5020I, A5039S, V5104L, T5220I, R5314M, and T5448I), eight mutations in the nsp3-Multi-domain (D1036E, L1249H, S1520F, F202526 P21V47, V204710, V2047, and T2648I), three mutations in nsp5-main proteinase (V3388I, N3405S, V3420L), three mutations in nsp14-Exonuclease (P6000S, T6249P, M6345V), 2 in nsp15-one (EndoRNAse) (K6486R and R6648G), 1 in nsp2 (T265I), one in nsp10-CysHis (R4387S), and 1 in nsp16 Methyltransferase (T7083I) (**Table S2**). We systematically performed the analysis predicting the deleterious effect of missense mutations in two genomes or more. Functional evaluation based on the SIFT (Sorting Intolerant From Tolerant) algorithm broke the tolerance index score of 15 missense mutations (found in two genomes or more), revealing its deleterious effect (**Table 1**).

**Table 1.** List of 15 non-synonymous amino acid substitutions with their tolerant effects.

| **Genes** | **AA position** | **Frequency (n=48)** | **Frequency (%)** | **Variation effect on protein** | **Tolerance index (SIFT)** |
| --- | --- | --- | --- | --- | --- |
| nsp2 | T265I | 12 | 11.11 | Benign/Tolerated | 0.62 |
| nsp3 | D1036E | 4 | 3.7 | Benign/Tolerated | 1 |
| nsp3 | V2047F | 5 | 4.63 | Deleterious | 0.04 |
| nsp3 | A2637V | 2 | 1.85 | Benign/Tolerated | 0.31 |
| nsp3 | T2648I | 5 | 4.63 | Deleterious | 0 |
| nsp12 | C4588F | 2 | 1.85 | Deleterious | 0.02 |
| nsp12 | T5020I | 9 | 8.33 | Deleterious | 0 |
| nsp12 | A5039S | 2 | 1.85 | Benign/Tolerated | 1 |
| nsp12 | V5104L | 4 | 3.7 | Deleterious | 0 |
| S | V6F | 5 | 4.63 | Benign/Tolerated | 0.15 |
| S | D614G | 48 | 44.44 | Benign/Tolerated | 0.62 |
| ORF3a | Q57H | 16 | 14.81 | Deleterious | 0 |
| ORF3a | L147F | 2 | 1.85 | Deleterious | 0.03 |
| N | R203K | 13 | 12.04 | Benign/Tolerated | 0.11 |
| N | G204R | 13 | 12.04 | Deleterious | 0.02 |

Eight missense mutations were found with deleterious effects, while the rest predicted to be benign/tolerated. Of the eight deleterious mutations, six were observed in the non-structural protein of the ORF1ab region; three (C4588F, T5020I, V5104L) could be deleterious for the nsp12-RdRp activity and two (V2047F, T2648I) in nsp3-Multi-domain. Likewise, two deleterious mutations (Q57H, L147) were found in the accessory protein ORF3a and one in the N protein (G204R).

### Haplotype network analysis

To estimate the number of introductions of the SARS-CoV-2 virus in Morocco, an analysis of the haplotype network was carried out using the 48 strains. Our results showed that these strains were grouped into five distinct clades, harboring 28 haplotypes from which three were dominant (**Figure 2**).

**Figure 2:**
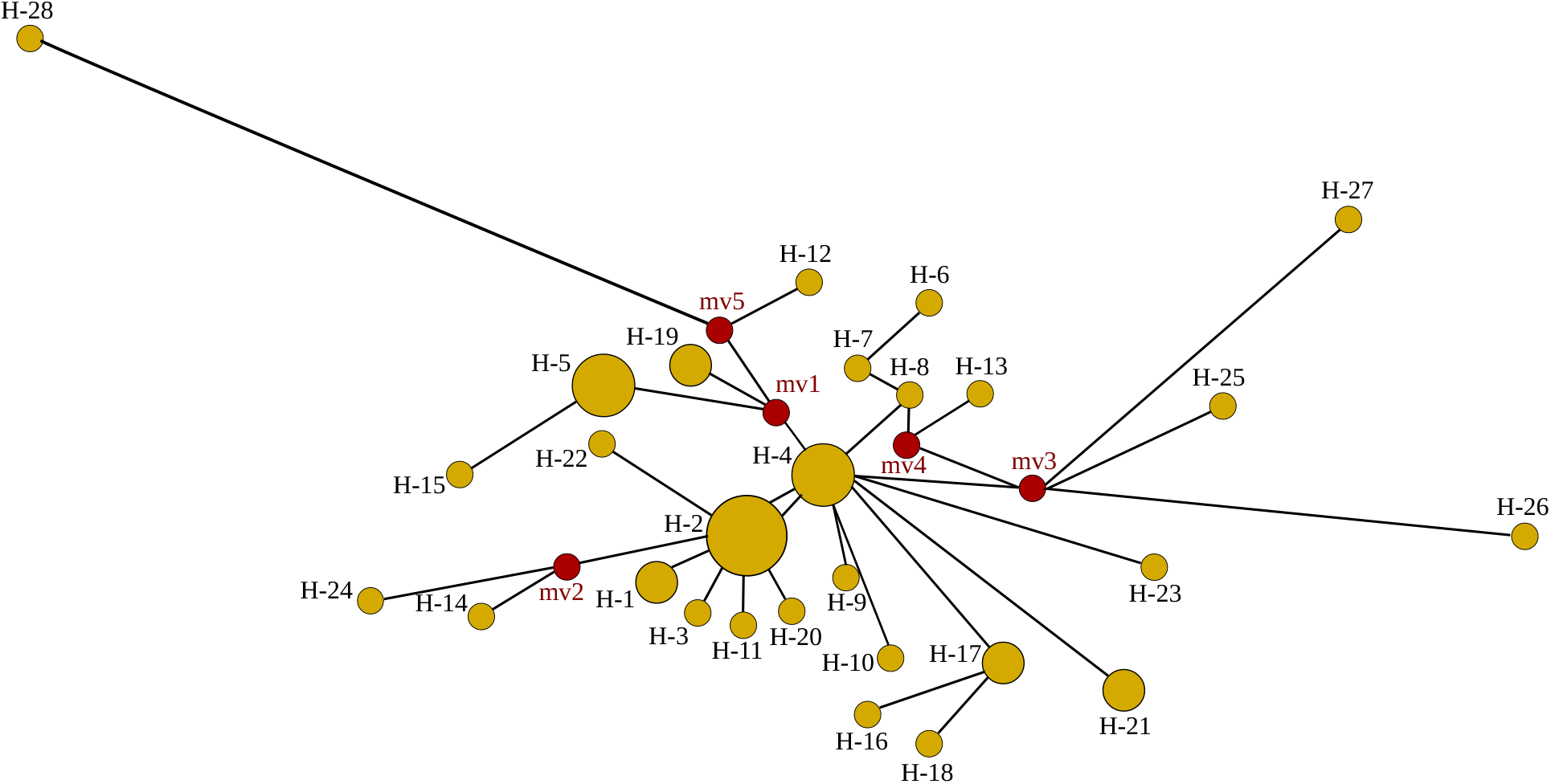
Haplotype network using genome-wide single-nucleotide variations (HN-GSNVs) of SARS-CoV-2 isolates from Morocco. Current samples have shown three major haplotypes harboring the majority of strains, H2 (11 strains), H4 (5 strains), H5 (4 strains), followed by H17, H21, and H1, indicating the predominance of H2 among Moroccan strains.

**In Figure 3**, the haplotypes are distributed in potential haplotype groups. Specifically, 43, 16, 47, and 88 haplotypes may be sub-haplotypes of primary ancestral haplotypes H1, H89, H25, and H59. Three of the 194 haplotypes that were found in Moroccan isolates were Moroccan-specific. The majority of Moroccan sequences were associated with haplotype group H25.

**Figure 3:**
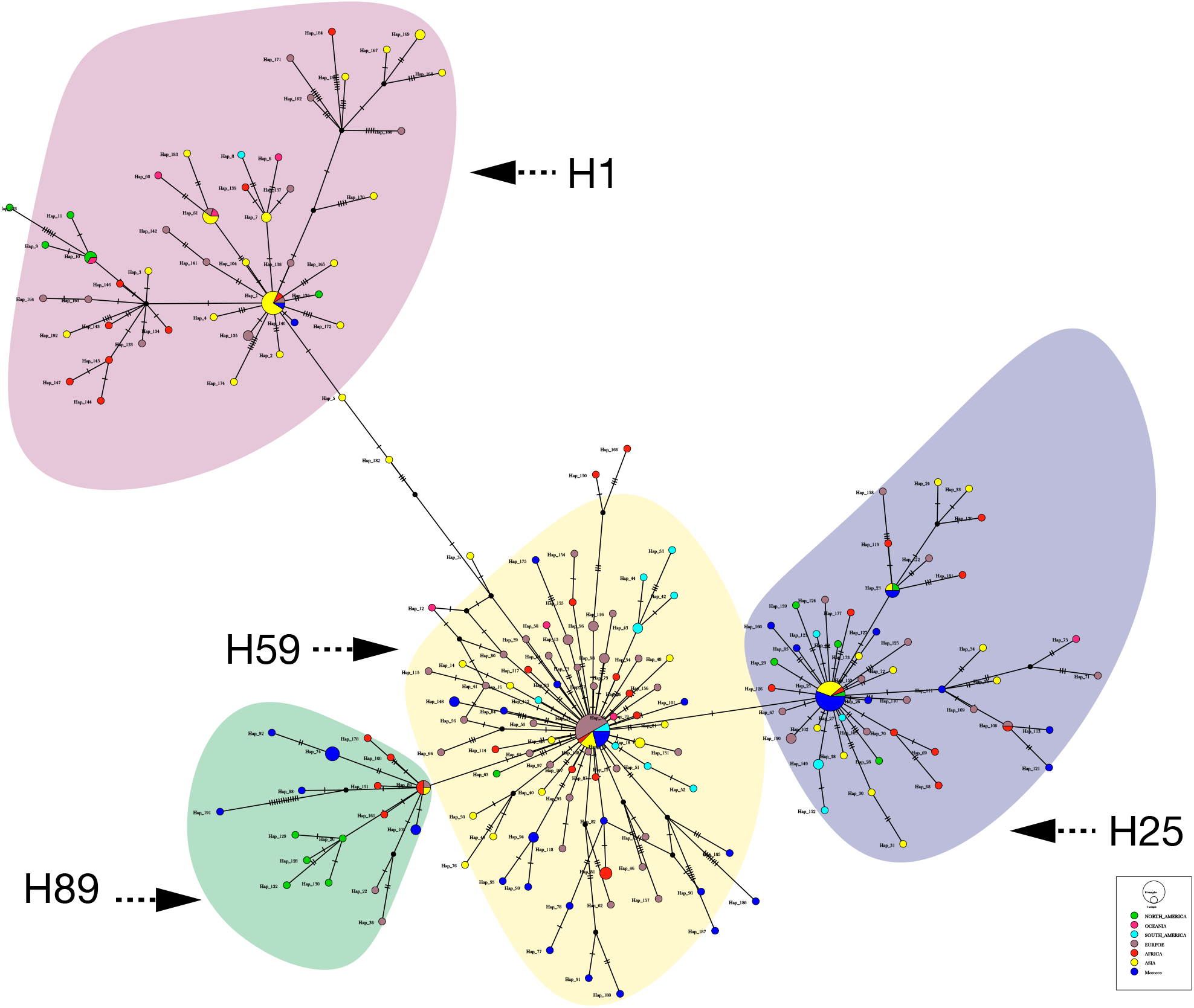
Haplotype network using genome-wide single-nucleotide variations (HN-GSNVs) of SARS-CoV-2 isolates in the world. Whole-genome sequences of SARS-CoV-2 isolates from 48 Moroccan patients were compared with 279 GISAID-available SARS-CoV-2 genomes from different parts of the world by median-joining SNV network analysis. Colors represent the sequence origin’s while the black strikes represent the number of mutations.

The haplotype groups’ distribution patterns differed in various geographic regions, with a few countries/territory specific. The 25 haplotypes harbored mainly strains from Asia, North American, and Africa, respectively, and are considered the ancestral strains. The number of haplotypes increased over time as new variants were continuously acquired in haplotype group 2, harboring the second biggest haplotype, including strains from Europe, Morocco, Asia, Africa, and South America, which originated 17 Moroccan haplotypes. H89 contained mainly strain from Asia, Africa, and Europe and give birth to five haplotypes from North America and Morocco. Lastly, group 4, which contains mostly strains from Asia, harbored fewer strains from Morocco.

### Phylogenetic analysis of Moroccan SARS-CoV-2 genomes with genomes belonging to six geographic areas

The phylogenetic analysis was carried out using 272 genomes from different countries belonging to the six continents (**Supplementary Material; Table S3**) to study the possible origin of SARS-CoV-2 strains circulating in Morocco. According to the GISAID nomenclature (**Figure 3**), our results revealed seven main clades: the “L” clade mainly contained genomes from Asia, while the other clades contained genomes belonging to different geographical areas.

**Figure 4:**
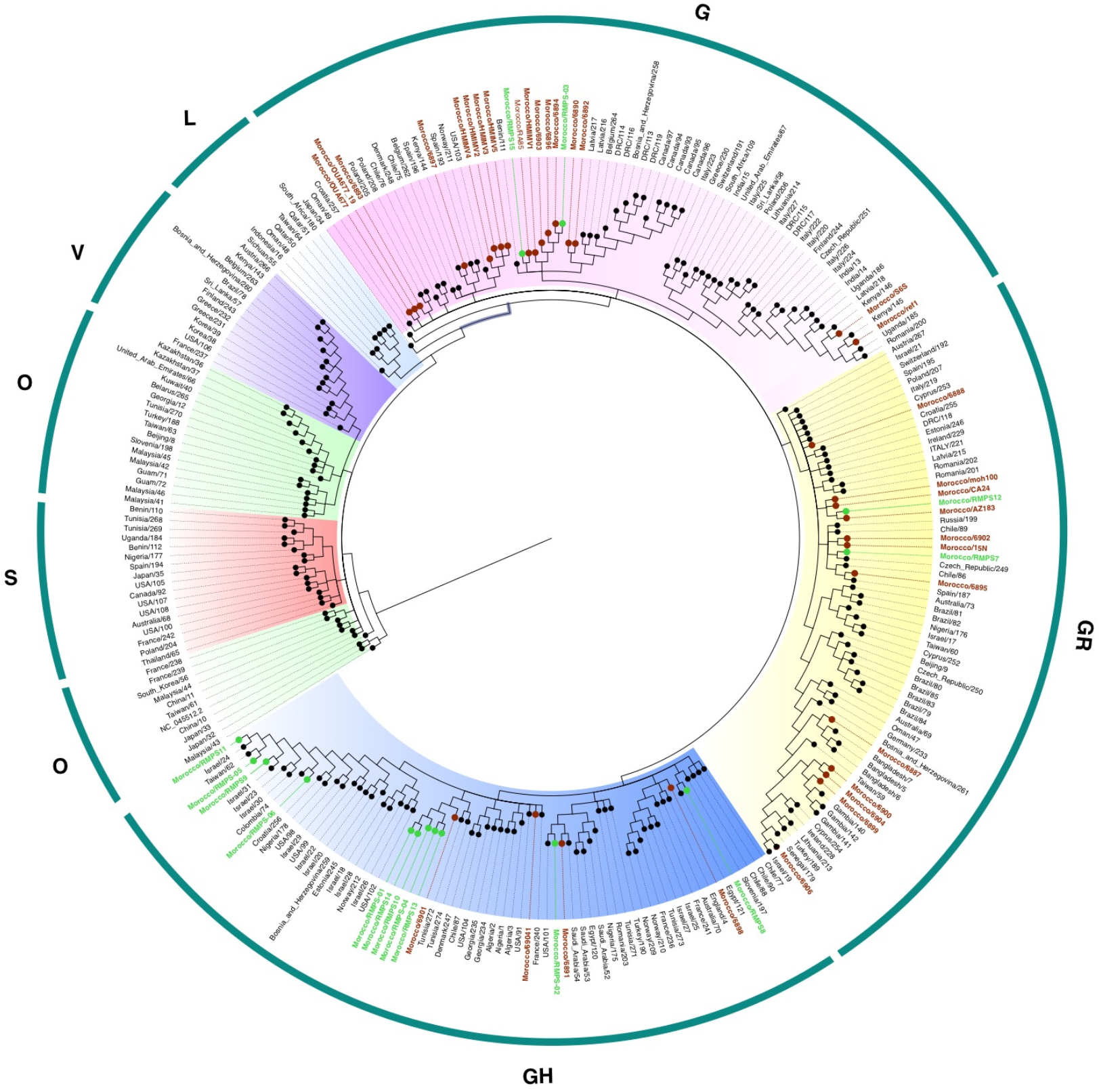
Phylogenetic tree based on 271 complete SARS-COV2 genomes from different geographic areas. The scale bar shows the branch’s length, which represents the change of nucleotides in the genome. The six Moroccan isolates newly sequenced in this study are represented by turquoise, and the other genomes from the same country (retrieved from the GISAID database), represented by red.

We observed that all Moroccan SARS-CoV-2 strains belonged to three close clades “G, GH, and GR,” harboring all the D614G mutation. These clades are also subdivided into several subclades. The G clade housed about half of the Moroccan strains and is mainly closely related to European origin strains, except three strains (Morocco / S6S, Morocco / ref1, Morocco / HMMV4) are closer to strains from Kenya and the USA. Meanwhile, among the Moroccan strains of the GH clade, five (Maroc / RMPS-01, Maroc / RMPS-14, Maroc / RMPS-04, Maroc / RMPS-10, Maroc / RMPS-13, Maroc / 6901) grouped close to the strains of Tunisia; In contrast, the other Moroccan strains of this same clade appear to share close sequence similarity with strains from different geographic areas, including Asia (Israel, Taiwan, Saudi Arabia), North America (USA), Europe (France, England), and South America (Colombia). On the other hand, the GR clade contained four Moroccan strains (Maroc / 6906, Maroc / 6899, Maroc / 6904, Maroc / 6900, Maroc / 6887) grouped with strains from the Gambia, Senegal (West Africa), and Bangladesh (Asia). Moreover, nine strains were grouped with European strains, mainly from Russia, the Czech Republic, Spain, and Cyprus. Overall, Moroccan strains were closely related to those from different continents, indicating different infection sources.

## Discussion

The monitoring of genetic variants plays a significant role in orienting the therapeutic approach for the development of candidate vaccines to limit the SARS-CoV-2 pandemic (19), as there is currently no proven effective treatment for SARS-CoV-2.

To date, the genetic diversity of strains of SARS-CoV-2 from Morocco is poorly documented. In this study, we performed a detailed analysis of genetic variants of forty-eight Moroccan strains, including nine newly sequenced, to provide new information on genetic diversity and transmission of SARS-CoV-2 in Morocco.

Genetic diversity could potentially increase the physical shape of the viral population and make it difficult to fight, or the opposite, make the virus weaker, which could be translated with a loss of its virulence and a decrease in the number of critical cases (20). Compared to the Wuhan-Hu-1/2019 reference sequence, Moroccan strains harbored 4 to 15 genetic variants per strain, of which 1 to 11 involve a change of amino acids. These results are consistent with the mutation rate previously reported in SASR-CoV-2 from different geographic areas (21-24), reporting a low frequency of recurrent mutations in thousands of SARS-CoV-2 genomes (21). In total, 108 variant sites have been identified, of which only 36% are present in two or more genomes. This result correlates with previous studies that SARS-CoV-2 evolved and diversified mainly by a random genetic drift, which plays a dominant role in propagating single mutations (25-27).

The mutations were distributed along the virus genome. The ORF1ab polyprotein region, known to be express 16 non-structural proteins (nsp1-nsp16), had several mutations that could affect their activity (3). Three non-synonymous deleterious mutations were found in the RdRp region (also called nsp12), a key part of the replication/transcription machinery, and which is proposed as a potential therapeutic target to inhibit viral infection (28, 29). Likewise, a deleterious mutation has been predicted in the NSP3 protein, In addition, two deleterious mutations have been observed in ORF3a, an accessory protein that makes it possible to regulate the interferon signaling pathway and the production of cytokines (30), and the structural N protein that plays a crucial function in the virus genome by regulating RNA transcription and modulating biological processes in infected cells.

RNA viruses acquire mutations readily, most of which are deleterious, and viruses carrying such mutations are eliminated. If a mutation reaches a high frequency, the mutation is expected to provide a selective advantage to the virus, usually manifested by a higher transmission efficiency. Three of these deleterious mutations (T5020I-nsp3, G204R-N, Q57H-ORF3a) have been previously reported (N-P) as hotspot mutations a large population belonging to different geographical areas.

Five non-synonymous mutations were common within at least two genomes, among them, D614G (in S protein) and Q57H (in OR3a). The D614G mutation is proximal to the S1 cleavage domain of advanced glycoprotein (31) and was of great interest due to their predominance in the six continents (32, 33). Alouane et al. (21) showed that this mutation appeared for the first time on January 24, 2020, in the Asian region (China); after a week, it was also observed in Europe (Germany). The Q57H mutation was taken away end of February in Africa (Senegal), Europe (France and Belgium), and North America (the USA and Canada). Likewise, our previous study (21) showed that D614G had no impact on the two-dimensional and three-dimensional advanced glycoprotein structures. Furthermore, four of these five mutations (D614G, Q57H, T265I, and T5020I) have been considered hotspot mutations in a large population (21, 22). With this in mind, we believe that this will not present a serious issue for vaccine development and that a universal candidate vaccine for all circulating strains of SARS-CoV-2 may be possible.

It should be noted that our predicted deleterious variants in protein structure lack experimental validation. Further exploration would be needed to confirm their potential effects further.

Our results of the haplotype network of Moroccan strains revealed five diverse clades with 28 haplotypes, indicating different infection sources. Besides the haplotype network, the phylogenetic analysis using a set of 273 strains representing the six continents revealed seven main clades. The most important clade contained approximately three-quarters of all the strains. All SARS-CoV-2 strains from North Africa harboring the D614G mutation belonged to this clade, except for three Tunisian strains. Interestingly, the Moroccan and Tunisian strains were closely related to those from Asia, Europe, South, and North America, which could indicate different sources of SARS-CoV-2 infection in these two countries.

## Conclusion

The results of this study provide valuable information on the diversity and impact of genetic variants of Moroccan strains of SARS-CoV-2 and their possible origins. This discovery could contribute to further in-depth investigations combining genomic data and clinical epidemiology of SARS-CoV-2 patients in Morocco, and suggest that, to date, the limited diversity seen in Moroccan SARS-CoV-2 should not preclude a single vaccine from providing global protection.

## Acknowledgments

We sincerely thank the authors and laboratories around the world who have sequenced and shared the full genome data for SARS-CoV-2 in the GISAID database. All data authors can be contacted directly via www.gisaid.org.

